# An ancestral role of pericentrin in centriole formation through SAS-6 recruitment

**DOI:** 10.1101/313494

**Authors:** Daisuke Ito, Sihem Zitouni, Swadhin Chandra Jana, Paulo Duarte, Jaroslaw Surkont, Zita Carvalho-Santos, José B. Pereira-Leal, Miguel Godinho Ferreira, Mónica Bettencourt-Dias

## Abstract

The centrosome is composed of two centrioles surrounded by a microtubule-nucleating pericentriolar matrix (PCM). Centrioles regulate matrix assembly. Here we ask whether the matrix also regulates centriole assembly. To define the interaction between the matrix and individual centriole components, we take advantage of a heterologous expression system using fission yeast. Importantly, its centrosome, the spindle pole body (SPB), has matrix but no centrioles. Surprisingly, we observed that the SPB can recruit several animal centriole components. Pcp1/pericentrin, a conserved matrix component that is often upregulated in cancer, recruits a critical centriole constituent, SAS-6. We further show that this novel interaction is conserved and important for centriole biogenesis and elongation in animals. We speculate that the Pcp1/pericentrin-SAS-6 interaction surface was conserved for one billion years of evolution after centriole loss in yeasts, due to its conserved binding to calmodulin. This study reveals an ancestral relationship between pericentrin and the centriole, where both regulate each other assembly, ensuring mutual localisation.

**Short summary:** The pericentriolar matrix (PCM) is not only important for microtubule-nucleation but also can regulate centriole biogenesis. Ito et al. reveal an ancestral interaction between the centriole protein SAS-6 and the PCM component pericentrin, which regulates centriole biogenesis and elongation.

## Introduction

The centrosome is the major microtubule organising centre found in animals, being composed of two microtubule-based cylinders, the centrioles, which are surrounded by an electron-dense proteinaceous matrix that nucleates microtubules, the PCM. The centriole can also function as a basal body, nucleating cilia formation. Centrioles are normally formed in coordination with the cell cycle, one new centriole forming close to each existing one, within a PCM cloud.

The first structure of the centriole to be assembled is the cartwheel, a ninefold symmetrical structure, composed of SAS-6, CEP135/Bld10, STIL/Ana2, amongst others (van Breugel et al., 2011; Kitagawa et al., 2011; Lin et al., 2013; Tang et al., 2011; Nakazawa et al., 2007). This is followed by centriole elongation through the deposition of centriolar microtubules which is dependent on components such as CPAP/SAS-4 (Tang et al., 2009; Schmidt et al., 2009; Kohlmaier et al., 2009). Remarkably, SAS-6 and CEP135/Bld10 can self-assemble *in vitro* to generate a ninefold symmetrical cartwheel (Guichard et al., 2017). *In vivo*, active PLK4 is known to recruit and phosphorylate STIL, which then recruits SAS-6 (Arquint et al., 2012, 2015; Moyer et al., 2015; Ohta et al., 2014). In a process called “centriole-to-centrosome conversion”, the centriole recruits centriole-PCM linkers, such as CEP192/SPD2 and CEP152/asterless, which contribute to both centriole biogenesis and PCM recruitment (Sonnen et al., 2013; Dzhindzhev et al., 2010; Hatch et al., 2010; Cizmecioglu et al., 2010). Finally, CDK5RAP2/CNN and pericentrin are recruited, which then recruit and activate γ-tubulin and create an environment favourable for concentrating tubulin, leading to the formation of a proficient matrix for microtubule nucleation and organization (Delaval and Doxsey, 2010; Megraw et al., 2011; Woodruff et al., 2017; Feng et al., 2017).

How do centrioles form close to each other? The cascade of phosphorylation and interaction events between centriole components leading to centriole biogenesis is an intricate succession of positive feedback interactions. That circuit leads to amplification of an original signal present at the centriole, such as the presence of active PLK4, or its substrate STIL, hence perpetuating centriole biogenesis there (Arquint and Nigg, 2016; Lopes et al., 2015). It is possible that centrosome function, including PCM recruitment and microtubule nucleation, additionally regulates centriole biogenesis, allowing for new centrioles to form close by. In support of this idea, when centrioles are eliminated in human cells by laser ablation, they form *de novo* within a PCM cloud (Khodjakov et al., 2002). Although several studies have unravelled centriole-PCM interactions, most focused on how the centriole recruits the PCM and not whether the PCM may regulate centriole biogenesis.

To define how the PCM regulates the localisation of individual centriole components while avoiding confounding effects of the many centriole-centriole component interactions that exist in animal cells, we established an heterologous system, taking advantage of the diverse centrosomes observed in nature. While centrioles are ancestral in eukaryotes, they were lost in several branches of the eukaryotic tree, concomitantly with the loss of the flagellar structure (Carvalho-Santos et al., 2010). Instead of a canonical centrosome, yeasts have a spindle pole body (SPB), a layered structure composed of a centriole-less scaffold that similarly recruits γ-tubulin and nucleates microtubules (Cavanaugh and Jaspersen, 2017). The timing and regulation of SPB biogenesis are similar to the one observed in animal centrosomes (Lim et al., 2009; Kilmartin, 2014; Ruthnick and Schiebel, 2016). It is likely that the animal centrosome and yeast SPB evolved from a common ancestral structure that had centrioles (Figure 1A), as early-diverged basal fungi such as chytrids (e.g. *Rhizophydium spherotheca*) have both a centriole-containing centrosome and flagella (Powell, 1980). By expressing animal centriole components in fission yeast we observed that the SPB recruits them. We further demonstrate that the conserved PCM component Pcp1/pericentrin recruits the centriole component SAS-6, revealing a yet unknown interaction in animals, which we show to be important for animal centriole biogenesis. Our work establishes a new heterologous expression system to study centrosome function and evolution and reveals an important role for pericentrin in recruiting centriole components and regulating centriole biogenesis and elongation in animals.

**Figure 1.**
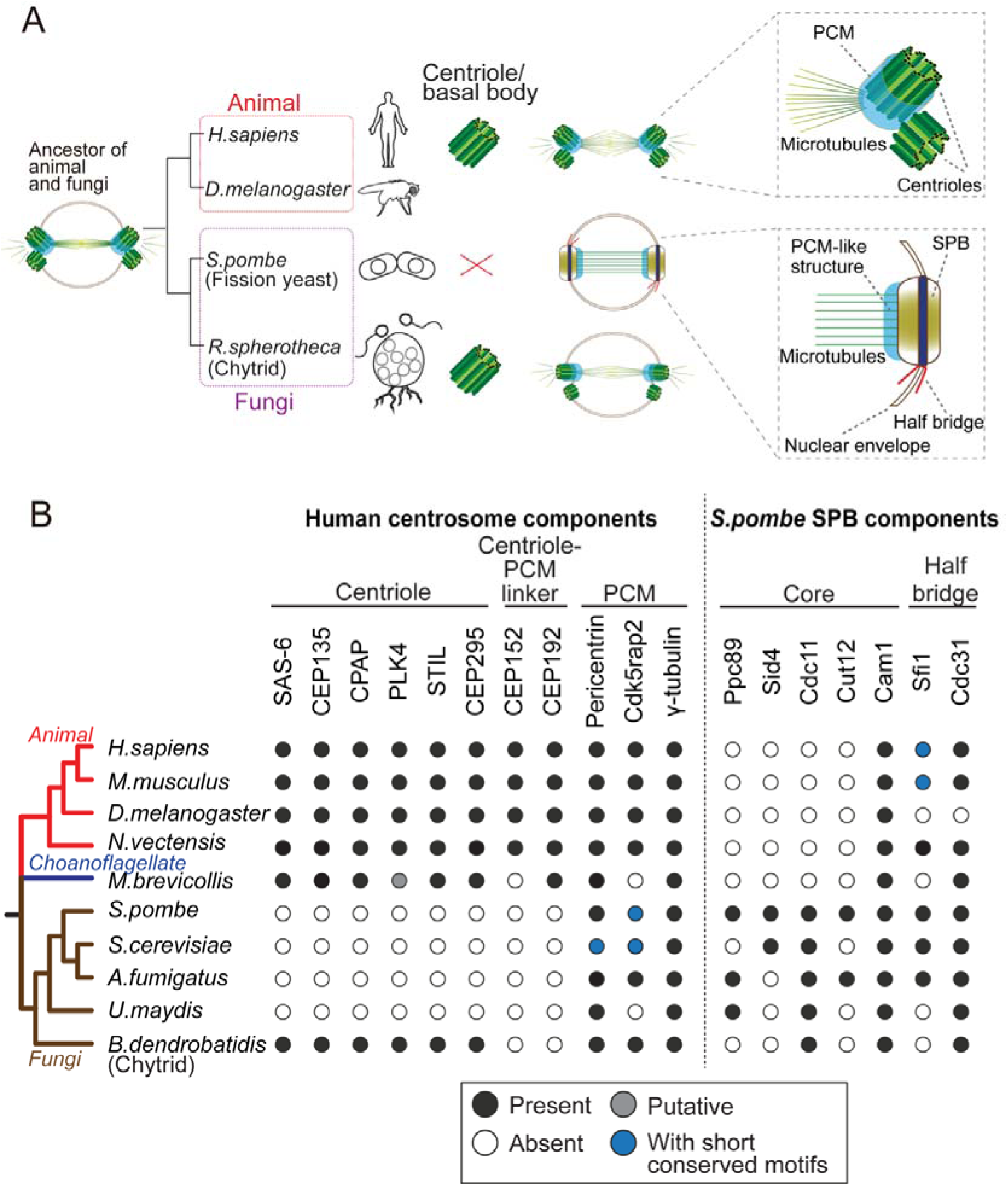
Evolution of the morphology and protein content of animal and fungi centrosomes. (A) The structure of the centrosome in mitosis in most animals, chytrids (flagellated fungi) and fission yeast. Animals and chytrids have a centriole/basal body and a canonical centrosome composed of a pair of centrioles surrounded by PCM, which anchors and nucleates microtubules. Fission yeast lacks a centriole but has a spindle pole body (SPB) inserted in the nuclear envelope. The SPB nucleates microtubules from the PCM-like structure, inside the nucleus. Parsimoniously, it is likely that the common ancestor of animals and fungi had a centriole-containing centrosome with a PCM structure (model shown). (B) Phylogenetic distribution of centrosome components in opisthokonts (animals, fungi and choanoflagellates). We searched for orthologues of components of the human centrosome localising to centrioles, centriole-PCM linkers and PCM, and the fission yeast SPB components. Black circles represent the presence of orthologues that were identified by the bidirectional best hit approach to the human or fission yeast proteins, respectively; grey circle represents the presence of putative orthologues identified by constructing phylogenetic trees; blue circles indicate that previous studies showed the presence of a protein with short conserved motifs (Kilmartin, 2003; Samejima et al., 2010; Lin et al., 2014) although we failed to identify it by the computational methods highlighted above; white circles indicate no detectable orthologue.

## Results

To establish yeasts as an heterologous expression system to study centrosome biogenesis, we analysed the conservation of the molecular components of the centrosome, searching for orthologs of the known animal proteins comprising: components of centrioles that are required for centriole biogenesis (SAS-6, STIL/Ana2, CPAP/SAS-4, CEP135/Bld10 and CEP295/Ana1), linkers of the centriole to the PCM, which are bound to the centriole and are required for PCM recruitment (CEP152 and CEP192), and the PCM itself, which is involved in γ-tubulin recruitment and anchoring (pericentrin, Cdk5rap2 and γ-tubulin itself). In addition, to better understand when the SPB originated, we also searched for orthologs of the fission yeast SPB components: the core scaffold proteins (Ppc89, Sid4, Cdc11, Cut12 and Cam1/calmodulin; Bestul et al., 2017; Chang and Gould, 2000; Krapp et al., 2001; Moser et al., 1997; Rosenberg et al., 2006) and the half-bridge proteins (Sfi1 and Cdc31/centrin; Kilmartin, 2003; Paoletti et al., 2003), which are required for SPB duplication.

Consistent with previous studies (Carvalho-Santos et al., 2010; Hodges et al., 2010), the proteins required for centriole biogenesis in animals were not identified in the fungal genomes, with exception of chytrids, which have centrioles (Figure 1B and Table S1). Centriole-PCM connectors were not found in both chytrids and yeasts. In contrast, when investigating the PCM composition, pericentrin and Cdk5rap2 were found in all fungal species, although the budding yeast Spc110 and Spc72 only share short conserved motifs with pericentrin and Cdk5rap2, respectively (Lin et al., 2014).

Regarding the SPB components, we found that Cam1 and Cdc31 are highly conserved across opisthokonts, which may reflect a conserved module or alternatively conservation related to MTOC-independent functions of these proteins. On the other hand, proteins such as Ppc89 and Sid4 are conserved only in yeasts but not in chytrids, suggesting that some of the yeast-specific structural building blocks of the SPB only appeared after branching into yeasts (Figure 1B and Table S1).

Altogether, these results suggest the yeast centrosome is very different from the animal canonical centrosome, not having centriole and centriole-PCM adaptors, and being composed of several yeast-specific SPB components. Our results suggest that different proteins are involved in assembling different modules of the centrosome, and that they can be lost when that module is lost, leading to divergence of the remaining structures. Importantly, while centriole components were lost and in turn SPB components are present, the PCM module seems to remain the same in animals and fungi in terms of composition and function, establishing fission yeast as an ideal system to study the interaction between individual centriole components and the PCM. We thus expressed key animal centriole components in fission yeast and asked whether they would interact with the PCM at the SPB or other yeast microtubule organising centres.

### Animal centriole components localise to the fission yeast SPB

We used individual *Drosophila* centriole components as they are well-characterized. We thus tested the localisation of *Drosophila* centriole components in fission yeast. We chose five critical components, SAS-6, Bld10/CEP135, SAS-4/CPAP, Ana2/STIL and PLK4 for the test. All the genes coding for these proteins are absent from the yeasts genomes (Figure 1B). Fission yeast SPBs are easily recognisable under light microscopy with fluorescent protein-tagged SPB marker proteins, such as Sid4 and Sfi1 (Kilmartin, 2003; Chang and Gould, 2000), which show distinct localisation as a clear focus. Therefore, we examined if animal centriole proteins can recognise and thus localise to the SPB.

GFP or YFP-tagged *Drosophila* centriole proteins were heterologously expressed under control of the constitutive *atb2* promoter (Matsuyama et al., 2008) or the inducible *nmt1* promoter (Maundrell, 1990) in fission yeast. We confirmed the expression of the fusion proteins with the expected sizes (Figure S1A). Despite the one billion years that separate yeasts from animals (Douzery et al., 2004; Parfrey et al., 2011), SAS-6-GFP, Bld10-GFP and YFP-SAS-4 co-localized with the fission yeast Sid4 to the SPB (Figure 2A). In addition to the localisation on SPB, YFP-SAS-4 signal was also weakly observed along the interphase cytoplasmic microtubules, likely reflecting its microtubule-binding capacity (Gopalakrishnan et al., 2011). In contrast, GFP-Plk4 and YFP-Ana2 did not localise to SPB and existed as aggregates in the cytoplasm (Figure 2A). We confirmed the expression of the fusion proteins with the expected sizes (Figure S1A), and the cells expressing centriole proteins which localise to the SPB grew as well as the control cells without any centriole protein (Figure S1B)

**Figure 2.**
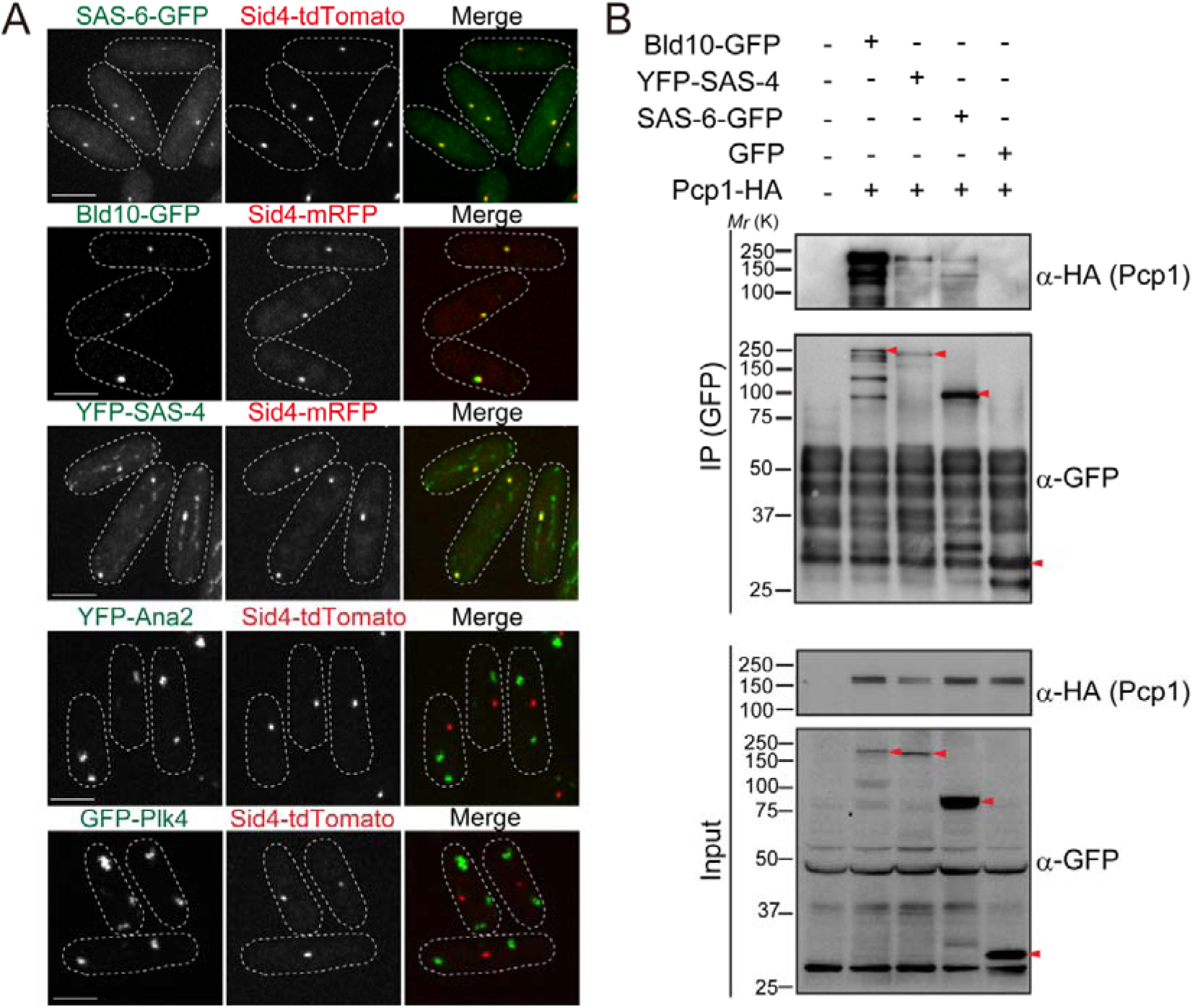
*Drosophila* centriole proteins localise to the centrosome of fission yeast. (A) SAS-6, Bld10 and SAS-4 localise to the SPBs but Ana2 and Plk4 do not. SAS6-GFP, Bld10-GFP and YFP-SAS-4 were expressed under control of the constitutive *atb2* promoter, and YFP-Ana2 and GFP-Plk4 under the *nmt1* promoter in fission yeast cells. (B) Physical interaction between the centriole proteins and fission yeast Pcp1. Protein extract was prepared from the fission yeast cells expressing HA-tagged Pcp1 and SAS-6-GFP, Bld10-GFP, YFP-SAS-4 or GFP. The GFP-tagged proteins were immunoprecipitated with anti-GFP antibody. Immunoprecipitates and inputs were analysed by western blotting using the indicated antibodies.

The result that SAS-6, Bld10 and SAS-4 independently localise to the SPBs, suggests that fission yeast SPB components can recruit them. These results were very surprising, indicating that SPBs and canonical centrosomes have diverged less than expected from their morphology or protein composition.

### Fission yeast pericentrin, Pcp1, interacts with the centriole proteins

We expected that the conserved PCM component localised on the SPB recruit the centriole components to the SPB. Then, we examined the interaction between the centriole proteins (SAS-6, Bld10 and SAS-4) and Pcp1, the fission yeast pericentrin ortholog, which recruits the γ-tubulin ring complex (γ-TuRC) to regulate mitotic spindle formation (Fong et al., 2010). In animals, pericentrin is a key component of the PCM, extending with its C-terminus at the centriole wall into the PCM (Lawo et al., 2012; Mennella et al., 2012). Indeed, we found that SAS-6, Bld10 and SAS-4 interacted with Pcp1 revealed by co-immunoprecipitation (Figure 2B). Amongst them, more Pcp1 was bound to Bld10 than the other two proteins, suggesting stronger affinity of Bld10 to the SPB.

### Fission yeast Pcp1 is required for SAS-6 recruitment

Next, we examined if Pcp1 is required for localisation of SAS-6, Bld10 and SAS-4 on SPBs using a temperature-sensitive mutant of Pcp1 (*pcp1-14*), in which the amount of Pcp1 protein is already reduced, and further reduced when grown at the restrictive temperature (Tang et al., 2014). These cells also arrest in mitosis when at the restrictive temperature. To compare the signal intensity in cells at the same cell cycle stage, mitosis, we introduced both in wild-type and *pcp1-14* background the mutation in a mitotic kinesin (*cut7-446)*, which fails in interdigitating the mitotic spindles and causes the cells to arrest in early mitosis (Hagan and Yanagida, 1990). Hereafter, we refer to the strains with *cut7-446* and *pcp1-14 cut7-446* alleles as the control and *pcp1* mutant, respectively. The intensity of SAS-6-GFP per SPB was significantly increased in the control cells when arrested in prometaphase (Figure 3A, D). We observed reduced SAS-6-GFP intensity in the *pcp1* mutant both at the permissive and restrictive temperatures (Figure 3A, D). This indicates that Pcp1 is required for SAS-6 recruitment to the SPB. Unlike SAS-6, the Bld10-GFP intensity was not reduced but slightly increased in the *pcp1* mutant (Figure 3B, E). It is possible that Bld10 is recruited to the SPB by another SPB component(s) and the possible recruiter might have been upregulated in the *pcp1* mutant. Although the intensity of SAS-4 was reduced in the *pcp1* mutant (Figure 3C, F), we found that the total protein of YFP-SAS-4 was lower in *pcp1* mutant while that of SAS-6-GFP and Bld10-GFP was comparable in control and *pcp1* mutant lysates (Figure 3G). We think that YFP-SAS-4 is stabilised by Pcp1 in the fission yeast cells, and collectively we cannot conclude that Pcp1 is required for YFP-SAS-4 localisation on SPBs. Since the localisation is clearly determined by Pcp1, we decided to explore further on how SAS-6 is recruited to the SPB by Pcp1.

**Figure 3.**
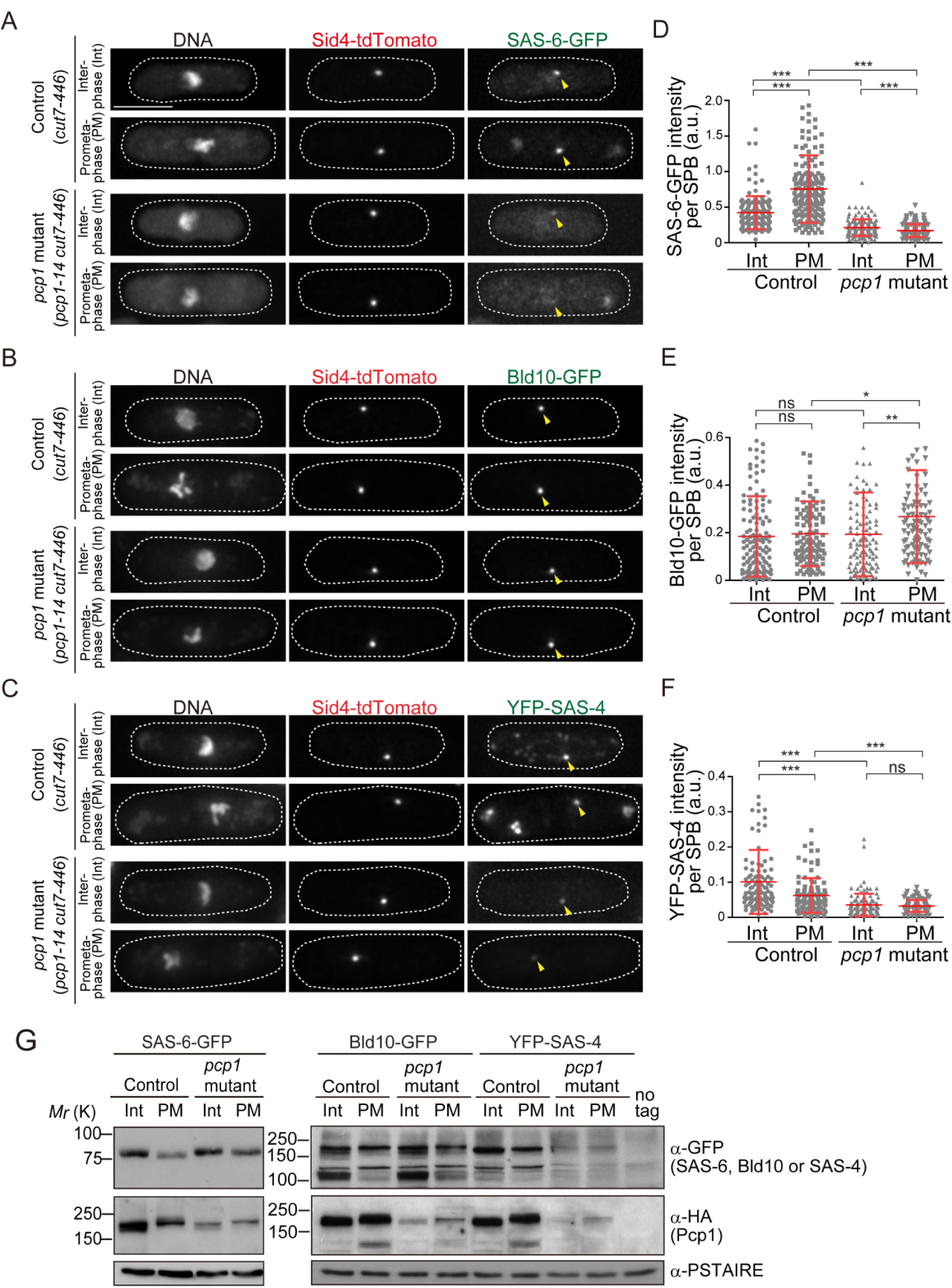
Fission yeast pericentrin-like protein Pcp1 is required to recruit SAS-6 to the SPB. (A-C) SAS-6-GFP, Bld10-GFP and YFP-SAS-4 intensities on the SPB in the *pcp1* mutant in asynchronous and prometaphase-arrested cells. The *cut7-446* (labelled “control”) and *cut7-446 pcp1-14* (labelled “*pcp1* mutant”) strains expressing SAS-6-GFP and Sid4-tdTomato were incubated at the restrictive temperature (36°C) for three hours to block cells in prometaphase due to the *cut7-446* mutation (temperature-sensitive allele of a mitotic kinesin, which causes failure in mitotic spindle formation.) Representative images of the cells collected before shifting the temperature (interphase, Int) and three hours after the shift to 36°C (prometaphase, PM) in control and *pcp1* mutant are shown. DNA was stained with DAPI. Arrowheads indicate the signal on SPB. Note that we also observed aggregation of SAS-6-GFP and YFP-SAS-4 in the cytoplasm both in the control and *pcp1* mutant at the restrictive temperature, which might be induced by the high-temperature stress. Scale bar, 5μm. (D-F) Quantification of the intensity of the centriole proteins per SPB in the indicated conditions. Means ± s.d. are shown in red (ns-not significant, * p<0.05, ** p<0.001, *** p<0.0001, Mann-Whitney U test). (G) Western blotting analysis of the protein extracts prepared from the indicated conditions.

It is known in *Drosophila* and human cells that phosphorylation and interaction of PLK4 with STIL/Ana2, facilitates STIL recruitment to the centriole and its interaction and recruitment of SAS-6 (Arquint et al., 2012; Ohta et al., 2014; Moyer et al., 2015; Vulprecht et al., 2012). However, neither Plk4 nor STIL are present in the fission yeast genome (Figure 1B and Table S1), suggesting that Pcp1/pericentrin is the additional molecular pathway for SAS-6 recruitment, and the ancestral interaction capacity is conserved in fission yeast.

### SAS-6 interacts with the conserved region of Pcp1, and Pcp1 is sufficient to recruit SAS-6

We reasoned that if the interaction between SAS-6 and Pcp1/pericentrin is an ancient and conserved connection, they should interact through an evolutionarily conserved domain in Pcp1. Subsequently, we determined which part of Pcp1 is required for its interaction with SAS-6. Full-length and truncation mutants of Pcp1 (N, M and C) were co-expressed with SAS-6-GFP (Figure 4A). Only the full-length and the C-terminal region containing the conserved PACT domain interacted with SAS-6 (Figure 4B). This region localises to the centriole wall in animals and is required for MTOC targeting both in animal and fungi (Gillingham and Munro, 2000).

We next asked whether Pcp1 could recruit SAS-6 ectopically. It is known that Pcp1 overexpression forms multiple Pcp1-containing foci in the cytoplasm (Flory et al., 2002). We thus examined if the cytoplasmic Pcp1 foci recruit SAS-6 to the ectopic sites (illustrated in Figure 4D). Overexpressed mCherry-tagged Pcp1 recruited SAS-6-GFP, but not GFP, to such foci (Figure 4D), indicating that Pcp1 can ectopically recruit SAS-6.

**Figure 4.**
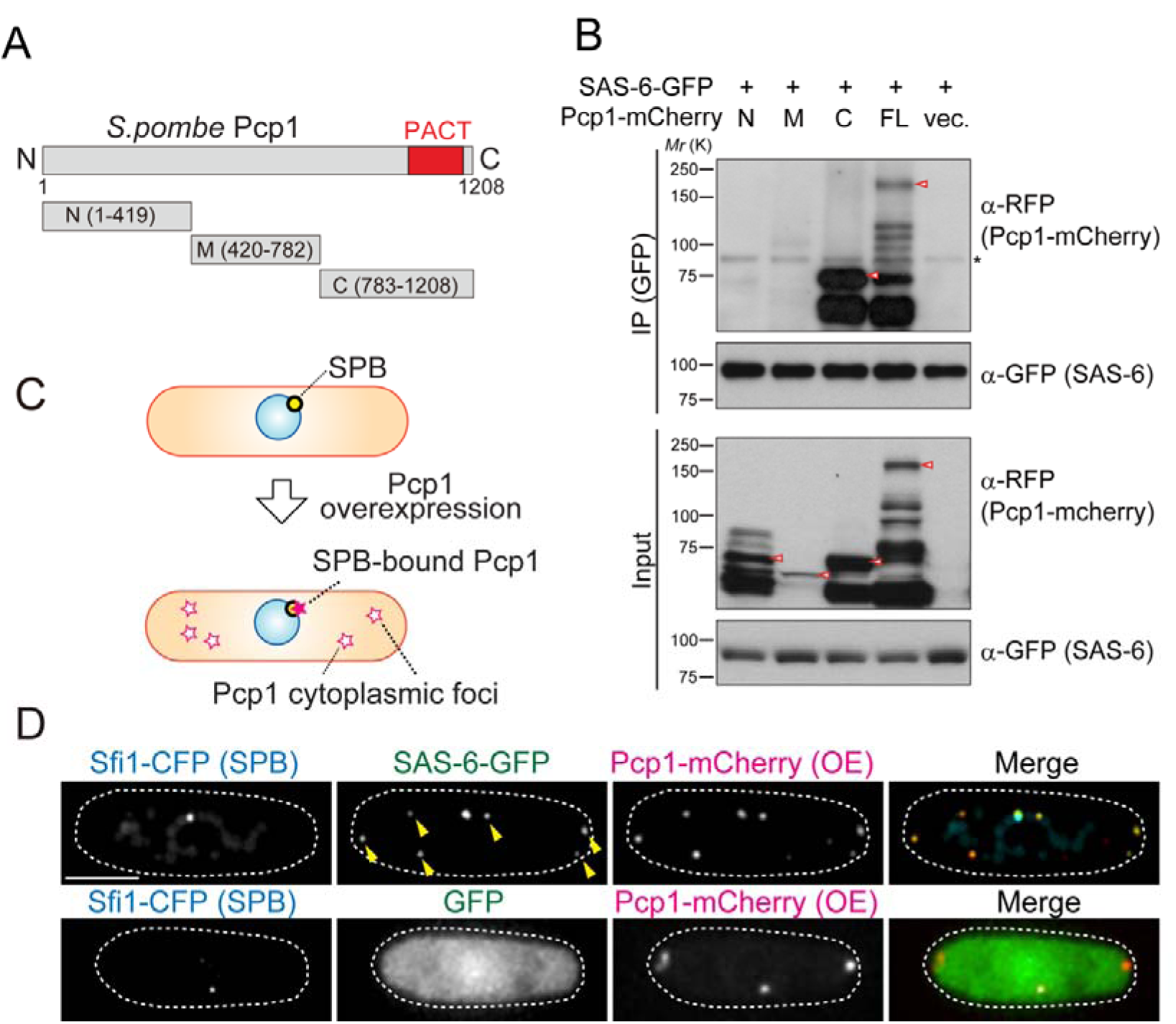
SAS-6 interacts with Pcp1 through the conserved carboxyl-terminal region, and Pcp1 is sufficient for SAS-6 localisation (A) Schematic illustration of the truncation constructs of Pcp1. (B) Pcp1 interacts with SAS-6 through the conserved carboxyl-terminal region. mCherry-tagged full-length or truncation mutants of Pcp1 were expressed in the cells constitutively expressing SAS-6-GFP. Immunoprecipitation was performed and analysed similarly as in Figure 2B. Red arrowheads indicate bands with the expected size of each fusion protein. The asterisk indicates non-specific bands. (C, D) Pcp1 is sufficient to recruit SAS-6. Overexpression of Pcp1 leads to the formation of Pcp1 containing cytoplasmic foci (schematic illustration, C). Pcp1-mCherry was overexpressed under control of *nmt41* promoter in the strain expressing SAS-6-GFP and GFP alone. Sfi1-CFP is shown (SPB marker). Arrowheads indicate the SAS-6-GFP signal on the Pcp1-cytoplasmic foci. Scale bar, 5μm.

### Conservation of the Pcp1/pericentrin - SAS-6 interaction

Our experiments suggest that there are conserved interactions between centrioles and PCM, whose evolution is constrained. They further suggest that PCM has an important role in recruiting centriole components. To test our prediction we examined in animal cells whether pericentrin interacts with SAS-6 and helps to recruit it to the centriole.

The pericentrin family varies in protein length but contains the conserved PACT domain in the C-terminal region (Figure 5A). Since we observed that fission yeast Pcp1 interacts with SAS-6 through the PACT-domain containing region, we tested whether SAS-6 interacts with the PACT domain of *Drosophila* PLP. EGFP-tagged SAS-6 and HA-tagged PLP fragment containing the conserved PACT domain were co-expressed in D.Mel cells. Consistent with the results obtained in fission yeast, SAS-6 interacts with PACT domain (Figure 5B). To verify whether this interaction is direct, we validated this result *in vitro*, by performing an *in vitro* binding assays using purified GST, GST-tagged SAS-6 and His-tagged PACT. GST-SAS-6 was specifically bound to His-PACT, indicating a direct interaction between these two proteins (Figure 5C).

**Figure 5.**
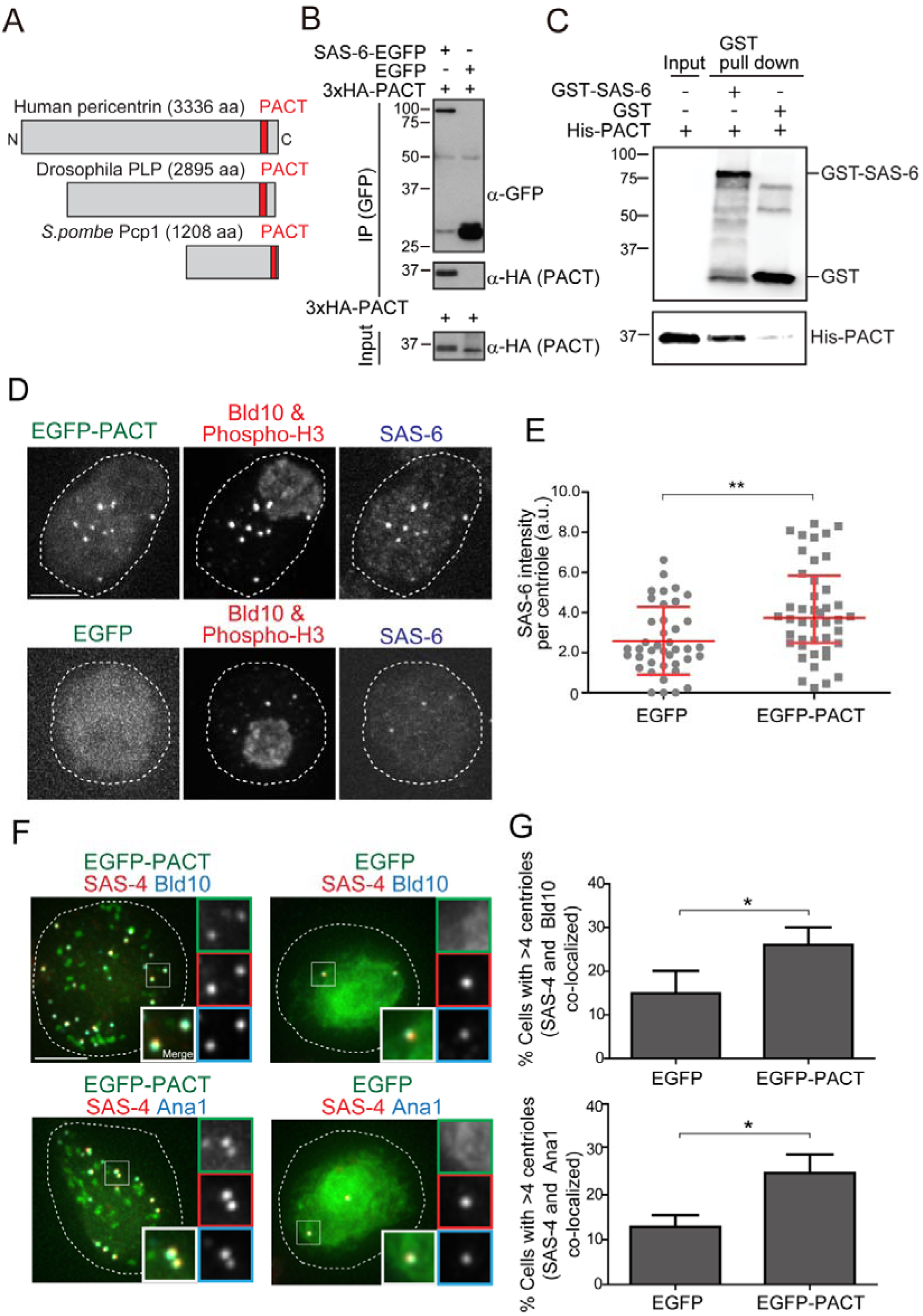
*Drosophila* pericentrin (PLP) conserved domain (PACT) interacts with SAS-6, and its overexpression causes centriole amplification. (A) Schematic illustration of human pericentrin, *Drosophila* PLP and *S.pombe* Pcp1. (B) Physical interaction between SAS-6-EGFP and the conserved *Drosophila* PACT domain. Protein extract was prepared from *Drosophila* tissue culture cells (D.Mel cells) expressing HA-tagged PACT, and SAS-6-EGFP or EGFP. The GFP-tagged proteins were immunoprecipitated with anti-GFP antibody. Immunoprecipitates and inputs were analysed by western blotting using the indicated antibodies. (C) Direct binding between SAS-6 and PACT. The *in vitro* binding assay was performed using purified GST or GST-fused SAS-6 and His-tagged PLP PACT. (D) Cells overexpressing EGFP-PACT or EGFP were arrested in mitosis by colchicine treatment for six hours and stained with the antibodies against Bld10 (centriole marker), phospho-H3 (mitotic marker) and SAS-6. (E) Quantification of SAS-6 intensity per centriole in cells overexpressing EGFP-PACT or EGFP arrested in mitosis. Means ± s.d. are shown in red (** p < 0.001, Mann-Whitney U test). Results are representative of three independent experiments (N >40 for each condition). (F) Cells overexpressing EGFP-PACT or EGFP were stained for two centriole markers (SAS-4 and Bld10, SAS-4 and Ana1) to count the centriole number. (G) Quantification of the cells with more than four centrioles (N>50, EGFP-positive cells). Data are the average of three experiments ± s.d. (*p < 0.05, Mann-Whitney U test). Note that although control *Drosophila* tissue culture cells already show cells with underduplicated and overduplicated centrioles as published before (Bettencourt-Dias et al., 2005), the expression of PACT leads to a significant amplification of centrioles.

### The conserved SAS-6-pericentrin interaction plays a role in centriole biogenesis

Previous evidence suggests that pericentrin may play a role in centriole biogenesis. Pericentrin is highly expressed and correlates with the levels of centrosome aberrations in acute myeloid leukaemia (AML) (Neben et al., 2004; Kramer et al., 2005). Moreover, overexpression of pericentrin in S-phase-arrested CHO cells induces the formation of numerous daughter centrioles (Loncarek et al., 2008). We wondered whether those effects were mediated by excessive SAS-6 recruitment since SAS-6 overexpression leads to supernumerary centriole formation (Peel et al., 2007; Strnad et al., 2007).

We first examined whether overexpression of EGFP-tagged PACT domain under actin5C promoter has an effect on centriole biogenesis. Cells overexpressing either EGFP-PACT or EGFP were arrested in mitosis with colchicine treatment in order to compare the intensity of SAS-6 signal at the centriole within the same cell cycle stage. Indeed, PACT-overexpressing cells increased the recruitment of SAS-6 per centriole, indicating that PACT recruits more SAS-6 (Figure 5D, E). This is consistent with our findings in fission yeast (Figure 4D). Since SAS-6 upregulation leads to centriole amplification (Leidel et al., 2005), we examined if PACT overexpression increases centriole number by staining with two combinations of reliable centriole markers (SAS-4 and Bld10, and SAS-4 and Ana1). Reflecting the increased recruitment of SAS-6 to the centrioles, we observed a significant increase in the percentage of cells with more than four centrioles compared to the EGFP control (Figure 5F, G).

### SAS-6 is recruited to the centriole by two complementary pathways

In animal cells, it is known that STIL/Ana2 recruits SAS-6 to the centriole (Arquint et al., 2012; Ohta et al., 2014; Moyer et al., 2015). However, it has been observed that STIL depletion does not completely prevent HsSAS-6 recruitment, and contribution of STIL does not fully account for centrosomal targeting of HsSAS-6 during interphase (Arquint et al., 2012; Keller et al., 2014). These results suggest that other factor(s) are required for centriole recruitment of SAS-6 in animals. Since overexpression of PACT recruits more SAS-6, we examined if PLP has a role in recruiting SAS-6 to the centriole in addition to STIL/Ana2. We performed a single depletion of PLP and Ana2 and a co-depletion of both together (Ana2 and PLP) in D.mel cells. We used *mCherry* depletion as a negative control for this experiment. Firstly, we investigated if SAS-6 recruitment is impaired by *Ana2* and *PLP* RNAi. We focused on mitotic cells to compare all cells at the same cell cycle stage. The intensity of SAS-6 per centrosome was quantified in each RNAi condition. Upon depletion of Ana2 or PLP, SAS-6 intensity per pole was significantly reduced compared to the control (Figure 6A, B). Moreover, double depletion of Ana2 and PLP caused an additive reduction of SAS-6 intensity. This result indicates that SAS-6 is recruited by two complementary pathways: Ana2 and PLP. Western blotting analysis confirmed that the targeted proteins were efficiently depleted, while total SAS-6 protein levels were comparable in all conditions (Figure 6C). We then asked whether altered recruitment of SAS-6 could affect centriole biogenesis. We observed that co-depletion of Ana2 and PLP increased the number of cells with (0-1) centriole indicating centriole biogenesis defects (Figure 6D, E). Importantly, the double depletion of Ana2 and PLP had a stronger effect on centriole biogenesis than the single ones, suggesting that both Ana2 and PLP contribute to the process by efficient SAS-6 recruitment.

**Figure 6.**
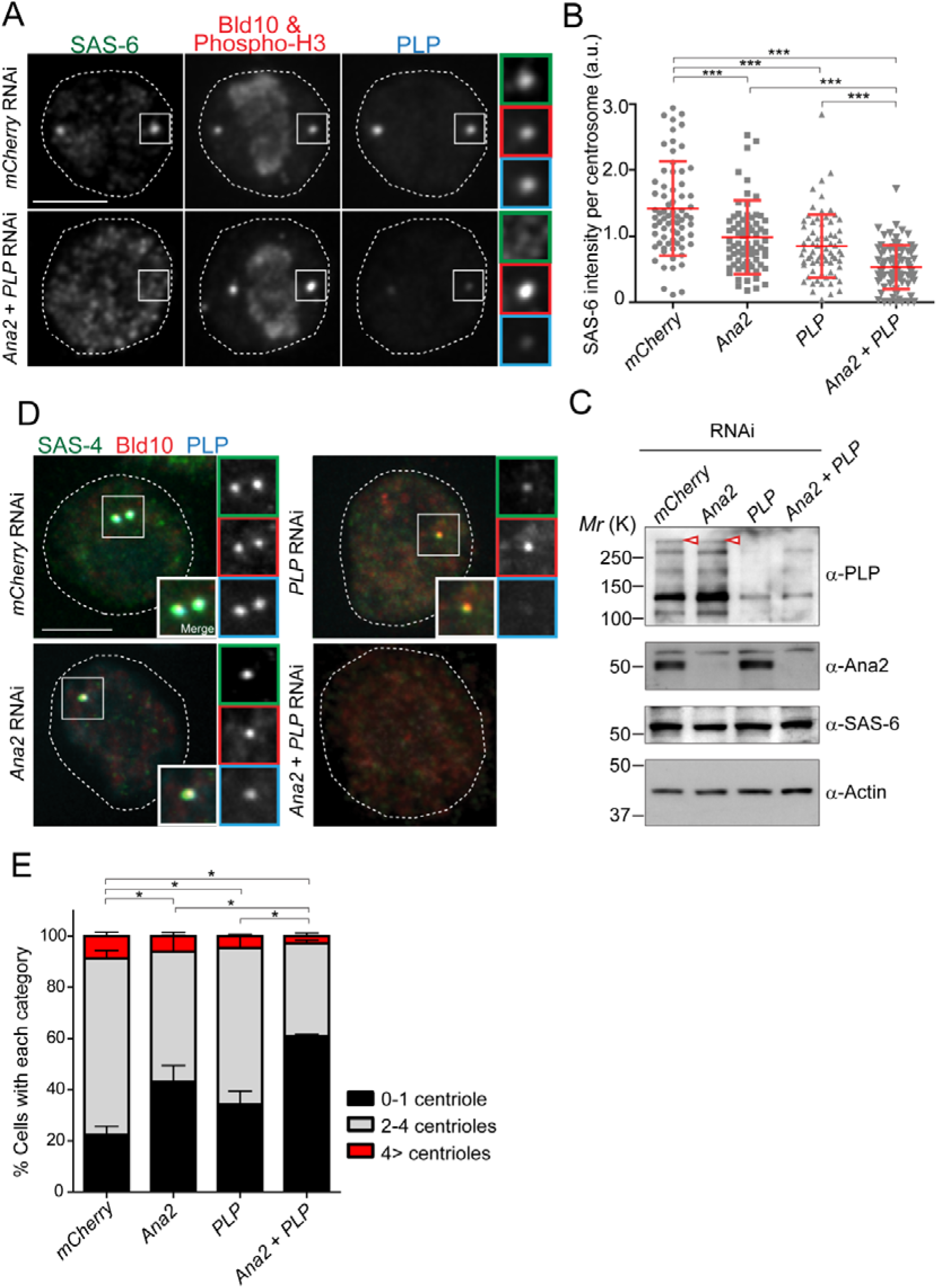
SAS-6-PACT complex formation complements the PLK4-STIL pathway, in recruiting SAS-6 to the centrosome and promoting centriole biogenesis. (A) Images of mitotic D.Mel cells after depletion of PLP and Ana2 by RNAi. D.Mel cells were depleted of PLP (*PLP* RNAi), Ana2 (*Ana2* RNAi), and were double-depleted with Ana2 and PLP (*Ana2* + *PLP* RNAi) (*mCherry* RNAi was used as negative control) for three days. Cells were immunostained with anti-SAS-6, Bld10, phospho-H3 and PLP antibodies. (B) Quantification of the SAS-6 intensity per centrosome in the mitotic cells in the indicated RNAi conditions. Means ± s.d are shown in red (*** p<0.001, Mann-Whitney U test). Results are representative of three independent experiments (N>50 for each condition). (C) Western blotting analysis of PLP, Ana2 and SAS-6 protein levels in the cells treated with the indicated dsRNAs using the antibodies against PLP, Ana2, SAS-6 and actin (loading control). Red arrowheads indicate expected bands of the longest PLP isoform. (D) Images of interphase cells after depletion of PLP and Ana2 by RNAi used for centriole counting. Cells were immunostained with anti-Bld10 (red), SAS-4 (another centriole marker, green), and PLP (cyan) antibodies. DNA was visualised with DAPI. (E) Quantification of centriole number per cell (N>100). Data are the average of three experiments ± s.d (* p<0.05, Mann-Whitney U test performed for the 0-1 centriole category).

### Drosophila pericentrin is required for SAS-6 recruitment and is important for centriole/basal body (BB) elongation *in vivo*

SAS-6 is required for centriole biogenesis (Rodrigues-Martins et al., 2007) and recently implicated in centriole elongation (Hilbert et al., 2016; Jana et al., 2018). Therefore, we wondered in addition to STIL/Ana2 if pericentrin is indeed required for SAS-6 recruitment to the centriole and thereby its assembly and maturation in a multicellular organism, such as *Drosophila*. We study this process during spermatogenesis, as centrioles are converted to basal bodies to form cilia, and elongate to more than 1 um, becoming highly visible and easy to study. To investigate this, we depleted PLP during centriole assembly and basal body maturation in spermatocytes (*PLPRNAi*) using *Gal4^Bam^* (Chen and McKearin, 2003; see timeline in Figure 7A). We studied its consequences in the localization of basal body components and basal body structure and, subsequently, in male fertility (Figure 7A). Unlike the *plp* mutant flies (Martinez-Campos et al., 2004; Galletta et al., 2014), the knockdown of PLP by RNAi did not affect centriole number in sperm cells (Figure 7B, C). Remarkably, the RNAi affected SAS-6 recruitment to the BBs (Figure 7D, E). Furthermore, this phenotype was associated with shorter BBs and male infertility (Figure 7E), a phenotype previously associated with reduced SAS-6 (Rodrigues-Martins et al, 2007; Jana et al, in press). Altogether, these results indicate that PLP is involved in sperm BB elongation by recruiting SAS-6.

**Figure 7.**
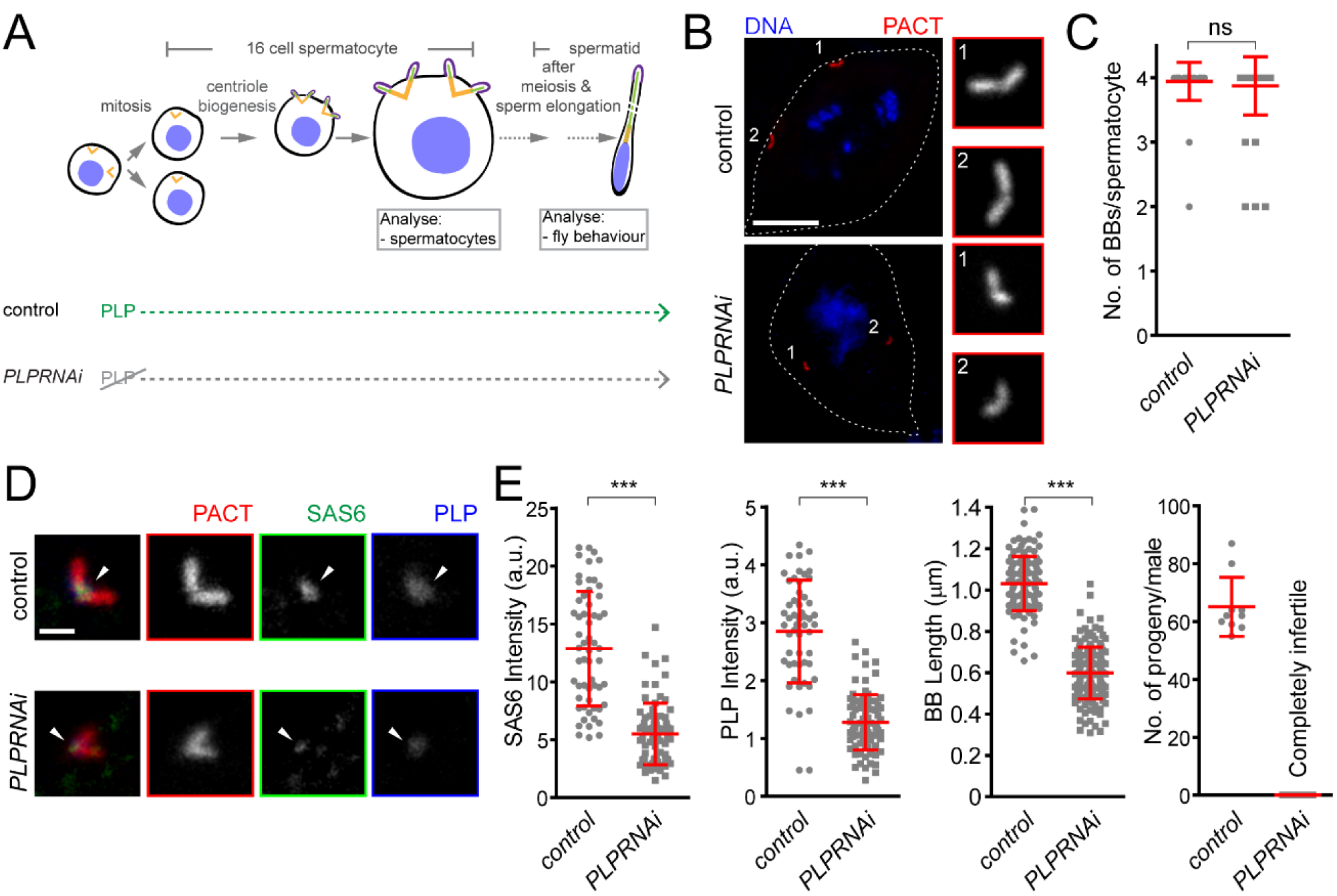
*Drosophila* pericentrin (PLP) is required for SAS-6 recruitment to the sperm centriole/basal body and for its elongation. (A) Schematic illustration of the experiments to deplete PLP during centriole biogenesis and elongation. (B) Representative images of mature spermatocytes in flies with different genotypes. PACT (red) is a commonly used marker for basal bodies (BB) and DAPI (blue) stains DNA. Insets show RFP::PACT close to the numbers (in grey scale). (C) Quantification of the number of BBs per cell in mature spermatocytes. (D) Representative images of mature spermatocyte BB in flies with different genotypes. RFP::PACT (red) marks BBs, Anti-SAS-6 (green) and Anti-PLP (blue) antibodies stain the proximal part of BB (arrowheads). (E) Quantification of SAS-6, PLP, BB length and the number of progeny in flies with different genotypes. We repeated all experiments three times. For SAS-6 and PLP intensities and BB length analysis, the number of BBs quantified for each genotype is N≥108 (54 pairs of BBs) and N≥128, respectively. The total number of males used for each histogram bar is N ≥10. Notably, given that moderate overexpression of PACT domain (using polyUbiquitin promoter) in the *plp* mutant fly fails to rescue the observed centriole as well as behaviour defects of the mutant (Martinez-Campos et al., 2004), we used RFP::PACT to study the sperm basal bodies in the knockdown experiments. Scale bars in (B) and (D) represent 10 and 1 μm, respectively. Means ± s.d are shown in red (ns-not significant, *** p<0.001, Mann-Whitney U test)

### The Calmodulin-PACT conserved interaction is likely to constrain the evolution of the PACT domain

We showed that the PACT domain has the capacity to interact with SAS-6 both in fission yeast and *Drosophila* cells, and the interaction between SAS-6 and pericentrin through PACT contributes to centriole biogenesis. It is possible that this interaction is ancestral, linking the two modules, the centriole and the PCM before the split between animals and fungi occurred one billion years ago. However, given the lack of centrioles and the divergence of the PCM in yeasts, we wondered why the PACT domain has retained the SAS-6 interaction surface. We hypothesised that other protein(s) interact with the same region of PACT domain as SAS-6, should be constraining the evolution of the interacting surface.

The PACT domain contains two highly conserved calmodulin (CaM)-binding domains (CBD1 and CBD2) (Galletta et al., 2014, Figure 8A). The interaction of pericentrin with calmodulin at CaM-binding domain is conserved and is important for its function both in animals and yeasts. In *Drosophila*, this interaction controls the targeting of PLP to the centrosome and is, therefore, critical for its function (Galletta et al., 2014). The interaction of the budding yeast pericentrin Spc110 with calmodulin is required for the SPB to nucleate microtubules and to form the spindle (Kilmartin and Goh, 1996; Spang et al., 1996; Stirling et al., 1994, 1996). Therefore, we asked whether SAS-6 interacts with this conserved segment covering CBD1 and CBD2 (hereafter called CBD). Similarly, as in Figure 5B, we co-expressed EGFP-tagged SAS-6 and the HA-tagged CBD in D.Mel cells to examine their interaction. Indeed, similarly as the PACT domain, CBD interacted with SAS-6-EGFP (Figure 8B). This result indicates that this conserved segment is sufficient for interaction with SAS-6.

**Figure 8.**
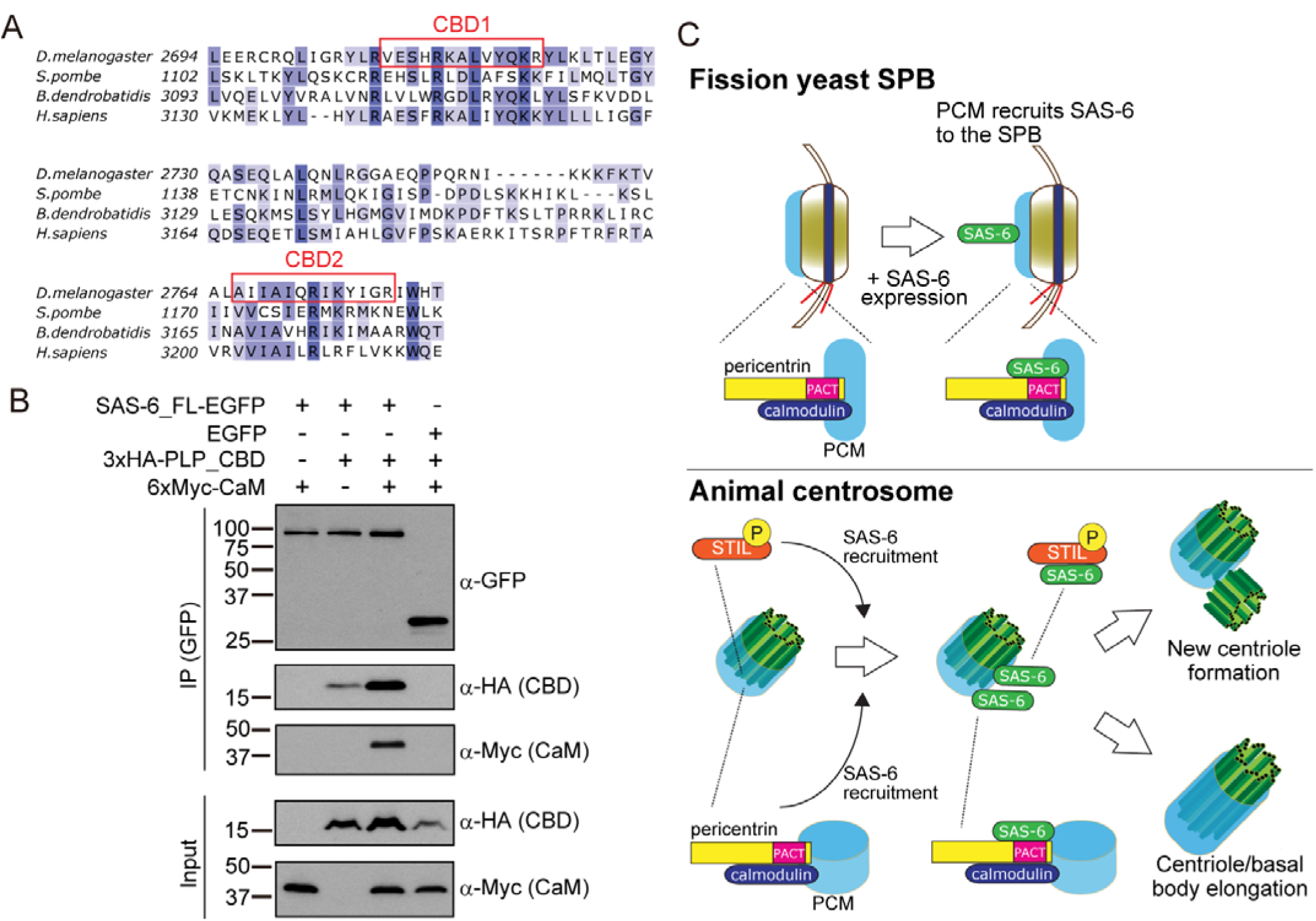
SAS-6 interacts with the calmodulin binding domain within PACT. (A) Graphical representations of the calmodulin (CaM)-binding domains (CBD1 and CBD2) within the multiple sequence alignment of the PACT domain of pericentrin proteins in the indicated species. The sequences of the PACT domain were aligned using Clustal Omega multiple sequence alignment tool and visually represented using Jalview software (McWilliam et al., 2013; Waterhouse et al., 2009). The alignments are color-coded in shades of blue for the percentage identity of amino acids; darkest blue (>80%), mid blue (>60%), light blue (>40%), white (<=40%). CBD1 and CBD2 are marked with red squares. (B) Complex formation between SAS-6, the highly conserved CBD within PACT domain and calmodulin. Protein extract was prepared from the D.Mel cells expressing SAS6-EGFP or EGFP, HA-tagged CBD and Myc-tagged calmodulin. The GFP-tagged proteins were immunoprecipitated with anti-GFP antibody. Immunoprecipitates and inputs were analysed by western blotting using the indicated antibodies. (C) Schematic representation of the ancestral role of PCM in the recruitment of centriole proteins, centriole biogenesis and elongation. When SAS-6 was heterologously expressed in fission yeast cells, it localised to the SPB through interaction with Pcp1/pericentrin (upper panel). This revealed a novel interaction between SAS-6 and pericentrin in animals, which is important for centriole biogenesis (lower panel). It is likely that SAS-6 is recruited to the pre-existing centriole prior to new centriole formation by two complementary pathways: PLK4-STIL/Ana2 and pericentrin. Even though yeast and animals are separated by one billion years of evolution, the pericentrin-SAS-6 interaction surface has been conserved likely because the binding of pericentrin to another centrosome component, calmodulin, has constrained its evolution.

Next, we asked whether SAS-6 is present in the same complex with calmodulin. Intriguingly, we found that SAS-6-EGFP, CBD and calmodulin formed a complex (Figure 8B). Since SAS-6-EGFP and calmodulin did not interact directly, we concluded that the complex forms through CBD. Moreover, we observed that the CBD polypeptide was stabilised in the presence of SAS-6-EGFP, and this was more pronounced with the addition of calmodulin, suggesting that formation of this complex might stabilise pericentrin (Figure 8B).

Our result suggests that the necessity for pericentrin-calmodulin interaction for both pericentrin function and stability, constrained the evolution of this segment of pericentrin, therefore retaining its affinity for SAS-6.

## Discussion

In this study, we examined the relationship between PCM and centriole components, taking advantage of a heterologous system, the fission yeast, which does not have centrioles. Surprisingly, the *Drosophila* core centriole components SAS-6, Bld10 and SAS-4 localised to the fission yeast SPB. In particular, SAS-6 was recruited to the SPB through interaction with the conserved PCM component, Pcp1/pericentrin, via its conserved calmodulin binding domain, within the PACT domain of Pcp1/pericentrin. Importantly, this interaction was also observed in animals (*Drosophila* cells). Further analysis revealed that pericentrin was required for SAS-6 recruitment to centrioles in addition to STIL/Ana2 pathway and for proper centriole biogenesis. It is estimated that animal and fungi diverged from the common ancestor about one billion years ago (Douzery et al., 2004; Parfrey et al., 2011). Our results reveal an ancestral relationship between the centriole scaffold and the PCM, in which PCM is needed for centriole component recruitment (Figure 8C). Therefore, the localisation of centriole birth is likely to be dictated by both positive regulatory feedbacks amongst centriole components, and also by the regulation of centriole component localisation by PCM.

### Implications of centriole biogenesis and elongation

SAS-6 is a critical structural component for centrioles. It assembles the cartwheel, which helps to define the centriole nine-fold symmetry (Kitagawa et al., 2011; van Breugel et al., 2011). It had been previously observed that the STIL/Ana2-dependent SAS-6 recruitment pathway, did not account for all SAS-6 present at the centriole (Arquint et al., 2012; Keller et al., 2014), but alternative pathways were not known. In this study, we serendipitously discovered that SAS-6 is recruited to the centriole through another new pathway mediated by pericentrin, though complementary to the previously characterized STIL/Ana2 pathway. Since human SAS-6 was previously seen to co-localize with pericentrin at the proximal end of the centriole where it accumulates from S phase (Keller et al., 2014), we suggest that pericentrin might recruit SAS-6 at that stage. Reflecting the role in the recruitment of SAS-6 to the centriole, we discovered that *Drosophila* pericentrin/PLP is required for centriole biogenesis in cultured cells, complementing the STIL/Ana2 pathway. Our data also provide a framework to understand why pericentrin overexpression leads to centrosome amplification (Kramer et al., 2005; Neben et al., 2004; Loncarek et al., 2008).

It is likely that the contribution of the pericentrin pathway is smaller than the STIL/Ana2 pathway and might only be observed after several cell cycles as it was not obvious before (Dobbelaere et al., 2008; Goshima et al., 2007; Martinez-Campos et al., 2004). Other cases have been described of cell type-specific differences observed for other centriole components, where phenotypes in tissue culture cells are enhanced (Franz et al., 2013; Delgehyr et al., 2012). Importantly, we also observed that pericentrin/PLP is required for SAS-6 recruitment to the basal bodies in testes and it is important for its elongation (Figure 7). A recent study also showed that the centriole length was reduced in wing disc cells in Drosophila *plp* mutant (Roque et al., 2018), suggesting an underappreciated generic role of PLP in centriole length control in the fly.

It seems that pericentrin/PLP-dependent SAS-6 recruitment is more important for centriole elongation, rather than biogenesis itself *in vivo*. Then, how are pericentrin and SAS-6 implicated in centriole elongation? The previous studies showed that SAS-6 is important for centriole maintenance (Izquierdo et al., 2014), and expression of the symmetry-changing SAS-6 mutant leads to reduced centriole length in human cells (Hilbert et al., 2016), suggesting that intactness of SAS-6 protein is indispensable for centriole stability and length control. A recent study suggested that *Drosophila* PLP helps to assemble electron-dense pericentriolar cloud-like structure around the mother centriole, which might be important for centriole length, cohesion and microtubule nucleation (Roque et al., 2018). We recently reported that SAS-6 not only constitutes the cartwheel structure but also is dynamically recruited to the spermatocyte BBs even after their cartwheel assembly. The later function of SAS-6 is required to elongate the BBs in the sperm cells (Jana et al., 2018). Here, our data suggest that pericentrin regulates the centriole length and stability by recruiting a part of the dynamic pool of SAS-6 to the centriole/BB.

We proposed that the binding of other molecules, including calmodulin, to the PACT domain of pericentrin, might constrain the evolution of that interacting surface, which also interacts with SAS-6. Calmodulin binding is important for pericentrin centriole-targeting, and also for its function and protein stability (Galletta et al., 2014). Moreover, the fission yeast *pcp1-14* mutant, which harbours a point mutation in the residue adjacent to CBD2, also exhibits reduced stability of the protein (Tang et al., 2014), implying a conserved role of calmodulin binding in stabilising Pcp1/PLP. Indeed, we observed that calmodulin binding stabilised PLP (Figure 7B). The previous study in budding yeast suggested that the calmodulin binding to the C-terminal region of Spc110 regulates the interaction between Spc110 and a critical inner plaque component Spc29 (Elliott et al., 1999). Perhaps calmodulin binding modulates the conformation of pericentrin into an “active form” which is stable and capable of interacting with centriole and SPB component(s), similarly to the regulation of calmodulin-dependent kinases by calmodulin (Crivici and Ikura, 1995).

### Implications for the study of cellular evolution and future application

Although fission yeast and other fungi lost centrioles, while acquiring the centrosome-equivalent SPBs, our study indicates that fission yeast SPBs still retain part of the ancestral PCM structure. Centrioles have a critical role in motility as basal bodies of cilia/flagella and in microtubule nucleation by recruiting the PCM. After centriole loss, the necessity to maintain microtubule nucleation, might have constrained the evolution of the PCM structure. Indeed, pericentrin orthologs in fungi, play essential roles in microtubule organization, mitotic spindle assembly, and cell cycle regulation similarly as in animals, and their function is dependent on calmodulin-binding (Geiser et al., 1993; Kilmartin et al., 1993; Flory et al., 2002; Fong et al., 2010; Chen et al., 2012). We think that the critical role of calmodulin-pericentrin interaction for proper microtubule nucleation has been retained in evolution, and constrained the evolution of pericentrin structure even after other interactors, such as SAS-6, were lost upon centriole loss.

It has been shown that many of the yeast genes can be substituted by human orthologs (e.g. rescuing the fission yeast *cdc2* mutant with the human CDK1 gene), indicating that the critical ancestral functions have been the same across a billion years (Lee and Nurse, 1987; Osborn and Miller, 2007; Kachroo et al., 2015). The present study demonstrated that the critical functional modules can retain not only the same function but also interaction capacities even when a binding partner is completely lost. The approach to heterologously expressing evolutionary-lost components of organelles in diverse organisms/cells could be useful to identify such novel interactions and important conserved interaction domains and divergent orthologue proteins across species.

Finally, since fission yeast SPB recruits centriole proteins, it might be feasible to use fission yeast to assemble multiple centriole components at SPBs. This synthetic biological approach could be useful to study the process of centriole biogenesis, such as the interaction between components and the order of recruitment, which will ultimately lead to the successful reconstitution of the evolutionary-lost centriole structure in fission yeast.

## Materials and methods

### Fission yeast strains and culture

The *S. pombe* strains used in this study are listed in Table S2. The strains were grown in yeast extract with supplements media (YE5S) or synthetic Edinburgh minimal media (EMM) in which ammonium chloride is replaced with glutamic acid (also called as PMG) with appropriate nutrient supplements as previously described (Moreno et al., 1991; Petersen and Russell, 2016).

### Plasmid DNA Constructions

Integration and expression vectors for *S.pombe* and *Drosophila* cells used in this study were constructed using the Gateway system (Invitrogen). All cDNA encoding *Drosophila* centriole components (SAS-6, Bld10, SAS-4, Ana2 and Plk4), PACT domain and calmodulin-binding domain of PACT (CBD) were amplified by PCR and cloned into pDONR221 vector.

To create integration plasmid for *S.pombe*, the pLYS1U-GFH21c (atb2 promoter, C-terminal GFP tag), pLYS1U-HFY1c (atb2 promoter, N-terminal YFP tag), pLYS1U-HFY1c (nmt1 promoter, N-terminal YFP tag), and pDUAL2-HFG1c (nmt1 promoter, N-terminal GFP tag) destination vectors (RIKEN Bioresource Center, Japan) were used. To express proteins in D.Mel cells, we used the destination vectors (*Drosophila* Genomics Resource Center, DGRC) containing the actin promoter termed: pAWG for the C-terminal EGFP tag, pAGW for N-terminal EGFP tag, pAHW for the N-terminal 3xHA tag, and pAMW for the N-terminal 6xMyc tag.

The plasmids for overexpression of fission yeast full length and truncated Pcp1-mCherry were constructed as follows. Each region of the *pcp1*+ gene was amplified by PCR and inserted into the SalI-NotI site of the expression plasmids pREP41 with mCherry at the carboxyl terminus in which gene expression is controlled under nmt1 promoter (Maundrell, 1990).

For recombinant protein expression, we used pGEX6p-1 (GE Healthcare) for GST and the Gateway pDEST15 (N-terminal GST, Thermo Fisher Scientific) destination vector for GST-DmSAS-6. To express 6xHis-PACT, the region of the PACT was amplified by PCR and inserted into the SalI-NotI site of the expression plasmid pET30-b (N-terminal 6xHis, Novagen). The plasmids used in this study are listed Table S3.

### Gene Targeting and Strain Construction in fission yeast

To generate the fission yeast strains expressing *Drosophila* centriole proteins SAS-6, Bld10 and SAS-4, the integration plasmids were linearized by digesting with NotI and integrated into each chromosomal locus. The integration into the targeted locus was verified by PCR.

### Drosophila Cell Culture and Transfections

*Drosophila* (D.Mel) cells were cultured in Express5 SFM (GIBCO) supplemented with 1× L-Glutamine-Penicillin-Streptomycin. DsRNA synthesis was performed as previously described (Bettencourt-Dias et al., 2004). Transient plasmid transfections were performed with Effectene reagent (QIAGEN) according to the manual. The primers used to amplify DNA templates for dsRNA synthesis are shown in Table S4.

### *Drosophila* stocks and culturing

All the fly stocks used in this study are described in Table S5 and publicly available stocks are listed in Flybase (www.flybase.org). Flies were reared according to standard procedures at 25 °C on corn meal media (Jana et al., 2016).

### Preparation of cell extracts, Western blotting and immunoprecipitation

*S.pombe* lysates were prepared using glass beads in extraction buffer (20 mM Hepes-NaOH (pH 7.5), 50 mM KOAc, 200 mM NaCl, 1 mM EDTA, 0.2% Triton X-100, and 0.1 mM NaF, additionally supplemented with 1× EDTA-free protease inhibitors (Roche) and 1 mM PMSF). Total cell lysates from D.Mel cells were prepared by resuspending cell pellets in lysis buffer described in (Galletta et al., 2014) (50 mM Tris (pH 7.2), 125 mM NaCl, 2 mM DTT, 0.1% Triton X-100, supplemented with 1× EDTA-free protease inhibitors (Roche) and 1 mM PMSF). Equal amounts of total cell lysates were separated in SDS-PAGE and analysed by immunoblotting. For the immunoprecipitation, the cell extract prepared from *S.pombe* or D.Mel cells were incubated for three hours at 4°C with Dynabeads Protein A (Thermo Fisher Scientific) pre-incubated with rabbit anti-GFP (Abcam). The beads were then washed three times with lysis buffer, and boiled in the Laemmli buffer for SDS-PAGE and western blotting.

### Protein purification and *in vitro* binding assay

*Escherichia coli* strain Rosetta (DE3) was transformed with the expression plasmid, and protein expression was induced at 25°C by the addition of 0.5 mM IPTG overnight. For purification of GST fusion proteins, cell pellets were lysed by sonication in lysis buffer (25 mM Tris pH 7.5, 150 mM NaCl, 1 mM EDTA, and protease inhibitors), and 10% Triton X-100 was added to the lysate at 0.5 % final concentration. GST fusion protein was affinity-purified with MagneGST Glutathione Particles (Promega). For purification of 6xHis-PACT, cell pellets were lysed by sonication in lysis buffer (50 mM NaH_2_PO_4_, 300 mM NaCl, 10 mM Imidazole, proteases inhibitors, and 0.1 % Tween, pH 8) supplemented with lysozyme at 1mg/ml final concentration. The protein 6xHis-PACT was affinity-purified with TALON^®^ Metal Affinity Resins (Clontech), and eluted with the elution buffer (50 mM NaH_2_PO_4_, 300 mM NaCl, 150 mM Imidazole, proteases inhibitors, 0.1% Tween) and then dialyzed was in the Dialysis Buffer (50 mM NaH_2_PO_4_, 200 mM NaCl, 5 mM Imidazole, pH8).

Recombinant GST or GST-DmSAS-6 beads were incubated with His-PACT supplemented with 500 μl of binding buffer (50 mM Na-HEPES, pH 7.5, 100 mM NaCl, 2 mM MgCl2, 1 mM DTT, 0.1% Triton X-100 and protease inhibitor) at 4°C for 1 hour, and the beads were washed three times in binding buffer. The proteins were eluted in SDS sample buffer and analysed by SDS-PAGE and western blotting.

### Immunostaining and imaging of *S.pombe* and D.Mel cells

Living or fixed *S.pombe* cells were mounted with a lectin-coated coverslip to immobilise before imaging. Immunostaining of D.Mel cells was performed as previously described (Cunha-Ferreira et al., 2013). Cells were mounted with Vectashield containing DAPI (Vector Laboratories). Cell imaging was performed on Nikon Eclipse Ti-E microscopes with Evolve 512 EMCCD camera (Photometrics) or DeltaVision Core system (Applied Precision) inverted microscope (Olympus, IX-71) with Cascade 2 EMCCD camera (Photometrics). Images were acquired as a Z-series (0.3 μm interval) and are presented as maximal intensity projections.

### Immunostaining, imaging and image analysis of Drosophila sperm cells

Testes from adult flies were dissected in testes buffer, transferred to poly-L-lysine glass slides, squashed, and snap frozen in liquid nitrogen as previously described (Jana et al., 2016). Then, testes were stained using different primary antibodies and secondary antibodies following the published method (Jana et al, 2016). Samples were mounted in Vectashield mounting media (Vector Laboratories) and they were examined in microscopes. Given that *Drosophila* has different stages of spermatocytes, we focused on the mature, large G2 spermatocytes and measured the total amount of SAS-6 and PLP at the basal bodies. All Confocal images were collected using Leica TCS SP 5X (Leica Microsystems, Germany) and processed in ImageJ.

### Male fertility tests

Fertility tests were performed by crossing single males with three wild-type females during 3 days. The progeny per tube was scored and averaged for ≥10 males for each genotype.

### Antibodies

We used the following primary antibodies for immunofluorescence (IF) or western blotting (WB): mouse anti-GFP (Roche, WB 1:1000), rabbit anti-GFP (Abcam, WB 1:1000), rat-anti-RFP (Chromotek, WB 1:1000), rat-anti-HA (Roche, WB 1:1000), rabbit-anti-Cdc2 PSTAIRE (Santa Cruz, WB 1:2000), rabbit-anti-Drosophila SAS-6 (gift from J.Gopalakrishnan, WB 1:500), rat-anti-Drosophila SAS-6 (gift from N.Dzhindzhev and D.Glover, IF 1:500), rat-anti-Ana2 (N.Dzhindzhev and D.Glover, WB 1:4000), rabbit-anti phosoho H3 (Millipore, IF 1:2000), rabbit-anti- Drosophila Bld10 (gift from T.Megraw, 1:5000), guinea pig-anti-Drosophila PLP (gift from G.Rogers, WB 1:1000), chicken-anti-Drosophila PLP (Bettencourt-Dias et al., 2005, IF 1:500), rabbit anti-Actin (Sigma, WB 1:2000), mouse anti-GST (Cell Signaling, WB 1:1000), mouse anti-His-tag (Novagen, WB 1:1000) and mouse anti-Myc (9E10) (Santa Cruz, WB 1:1000).

The following secondary antibodies were used: DyLight 488 donkey anti-Rat IgG (Bethyl Laboratories, 1:100), Rhodamine Red Donkey anti-rabbit IgG, Cy5 Donkey anti-chicken IgY, Cy5 Donkey Anti-rat IgG (Jackson ImmunoResearch, 1:100) for IF; horseradish peroxidase (HRP)-conjugated donkey anti-mouse IgG, HRP donkey anti-rabbit IgG, HRP donkey anti-guinea pig IgG (Jackson ImmunoResearch, 1:5000), HRP goat anti-rat IgG (Bethyl Laboratories, 1:5000), IRDye 800CW goat anti-mouse IgG, IRDye 680CW goat anti-Rabbit IgG (LI-COR, 1:10000), for WB.

### Fluorescence intensity quantification and centriole counting

For fluorescence intensity quantification, Z-stack images (0.3 μm intervals) of 21 sections (for fission yeast) or 61 sections (for D.Mel cells) were acquired. Integrated intensity of the fission yeast SPB (6×6 pixel) or the centrosome/centriole in D.Mel cells (7×7 pixel) were recorded in the maximum projected images. Cellular background intensity was subtracted from the SPB or centriole intensity. Image processing and quantification were performed in ImageJ.

### Statistical Analysis

The statistical analysis (non-parametric Mann Whitney U test) was performed in Graphpad Prism version 5.0 software.

### Bioinformatics analysis to predict orthologs

Complete proteomes were downloaded from Ensembl and Ensembl Genomes databases. EST information was downloaded from JGI. We performed orthology prediction based on multiple methods. Candidate orthologs were identified using pairwise sequence-based (BLASTP and phmmer) and domain-based (hmmsearch) methods (Altschul et al., 1990; Finn et al., 2015). The orthologs were then manually verified by (1) confirming the presence of critical domains and motifs, (2) using the bidirectional best hit approach. If results were ambiguous (i.e. there are multiple candidates), a putative ortholog was identified by constructing phylogenetic trees of the protein family using MrBayes 3.2.5 (Ronquist and Huelsenbeck, 2003) The species tree was downloaded from NCBI taxonomy.

## Acknowledgements

The authors thank E. Levy and P. Beltrão for critical reading of the manuscript, I. Hagan and P. Tran for helpful discussion. We are grateful to M.Gomes and C. Bicho for help with experiments. We thank I. Hagan, M. Sato, T. Toda, K. Tanaka, JQ. Wu, A. Paoletti, K. Gould, T. Matsumoto, M. Bornens, T. Megraw, N. Dzhindzhev, D. Glover, G. Rogers, B. Galletta, N. Rusan, RIKEN BRC and NBRP (Japan) for yeast strains, antibodies and plasmids. D.Ito is supported by a long-term fellowship from the Human Frontier Science Program (LT000344/2013), and postdoctoral fellowships from the Uehara Memorial Foundation, Japan and Fundação para a Ciência e a Tecnologia (FCT) (195/BPD/17). Work in the laboratory (CCR) of M.Bettencourt-Dias is funded by the Gulbenkian Foundation, a European Research Council Consolidator Grant (CoG683528), and FCT grant (PTDC/BIM-ONC/6858/2014).

---

**Figure S1 (related to Figure 2).**
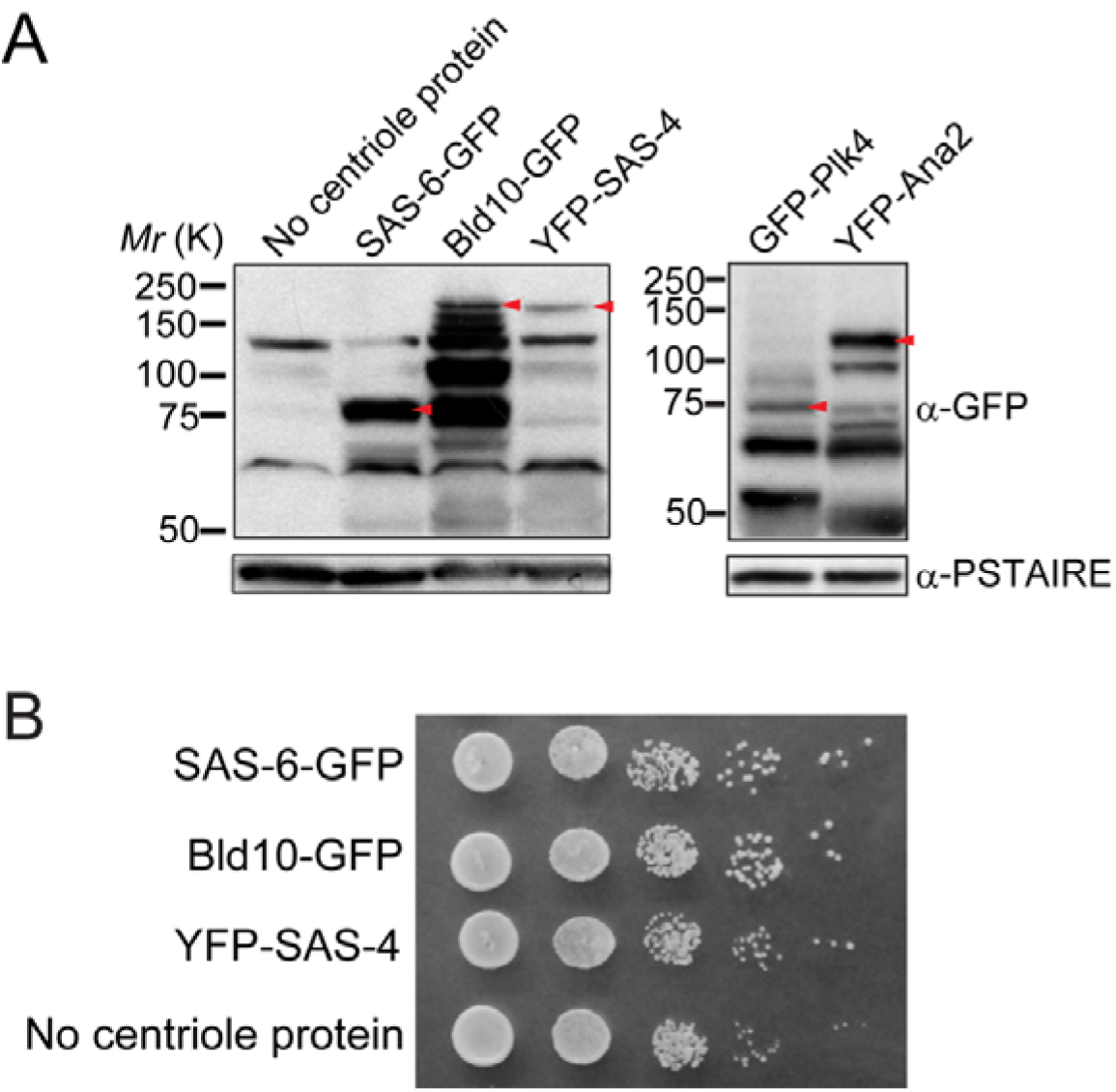
Expression of *Drosophila* centriole proteins in fission yeast and the growth of fission yeast strains. (A) Western blotting analysis of the protein extracts prepared from the strains used in Figures 2A. Each red arrowhead indicates the bands with the expected size of the fusion protein. (B) Serial dilution assay of the cells expressing indicated *Drosophila* centriole components under control of the constitutive *atb2* promoter, which are shown in Figure 2A. Note that all the strains expressing the centriole protein grew similarly as the control (no centriole protein).

**Table S1 (Related to Figure 1).**
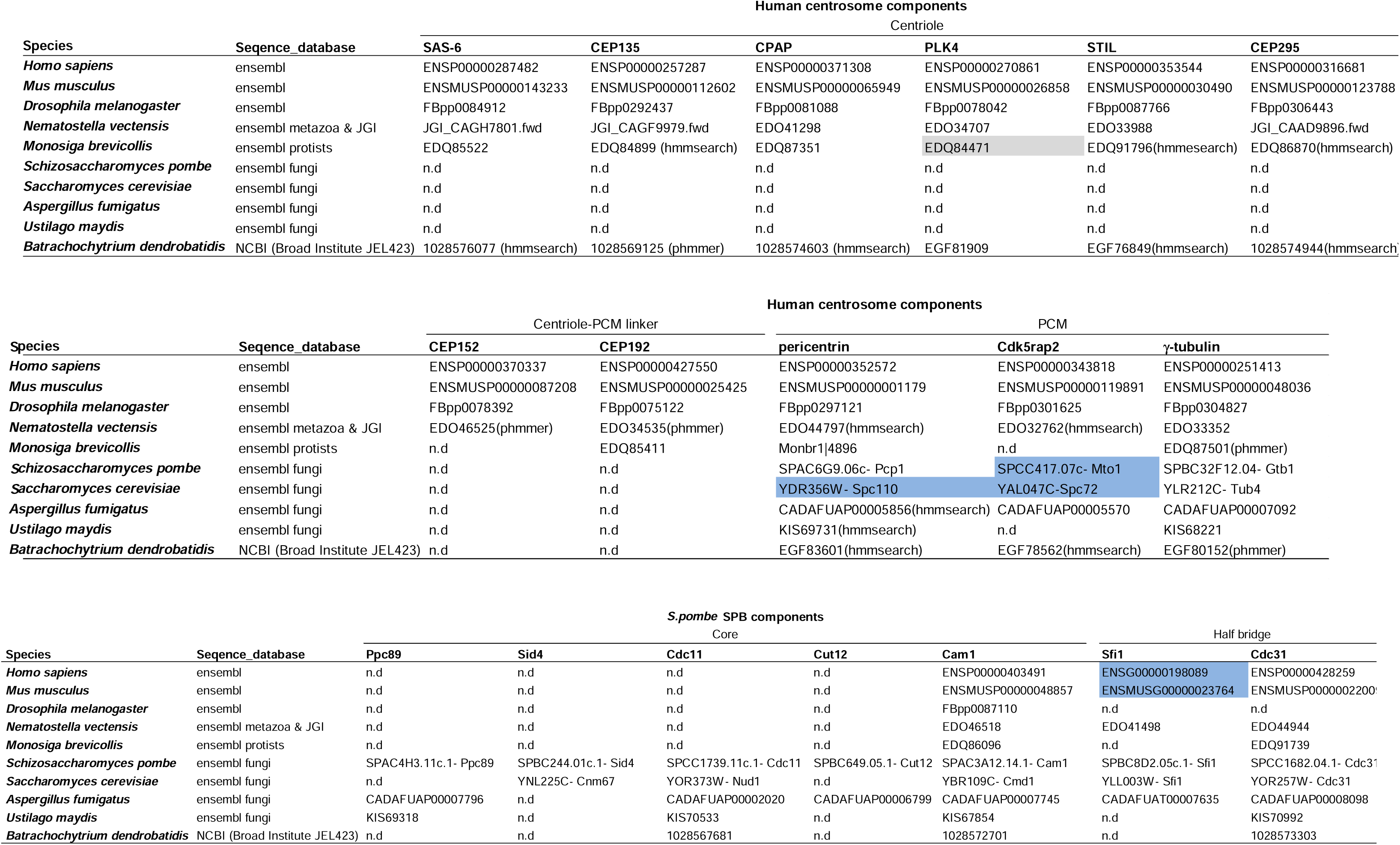
List of the predicted orthologs of the human centrosome and *S.pombe* SPB components in animal and fungi species. The gene IDs of the orthologs are shown if identified by the bidirectional best hit approach (see Figure 1 and the Materials and Methods for detail). Grey box represents the putative orthologues identified by constructing phylogenetic trees; blue box indicate the protein with short conserved motifs; n.d stands for “Not detected”. When predicted by hmmsearch or phmmer, it is indicated in the parenthesis.

**Table S2.** Fission yeast strains used in this study

|  |  |  |
| --- | --- | --- |
| Figure 2A | DI 456 | *h- leu1-32 ura4-D18 lys1::P<sup>atb2</sup>-DmSAS6-GFP-FLAG-6xHis-ura4+ sid4-tdTomato-natMX6* |
|  | DI 190 | *h- leu1-32 ura4-D18 lys1::P<sup>atb2</sup>-DmBld10-GFP-FLAG-6xHis-ura4+ sid4-mRFP-natMX6* |
|  | DI 202 | *h- leu1-32 ura4-D18 lys1::P<sup>atb2</sup>-6xHis-FLAG-YFP-DmSAS4-ura4+ sid4-mRFP-natMX6* |
|  | DI 655 | *h- ura4-D18 lys1::nmt1-6xHis-FLAG-YFP-DmAna2-ura4+ sid4-tdtomato-natMX6* |
|  | DI 665 | *h+ ura4-D18 leu1::nmt41-6xHis-FLAG-GFP-DmPlk4-ura4+ sid4-tdtomato-natMX6* |
| Figure 2B | DI 492 | *h- leu1-32 ura4-D18 lys1::P<sup>atb2</sup>-DmSAS6-GFP-FLAG-6xHis-ura4+ sid4-tdtomato-natMX6 pcp1-HA-hphMX6 cut7-446* |
|  | DI 706 | *h+ leu1-32 ura4-D18 lys1::P<sup>atb2</sup>-DmBld10-GFP-FLAG-6xHis-ura4+ sid4-tdtomato-natMX6 pcp1-HA-hphMX6 cut7-446* |
|  | DI 709 | *h+ leu1-32 ura4-D18 lys1::P<sup>atb2</sup>-6xHis-FLAG-YFP-DmSAS4-ura4+ sid4-tdtomato-natMX6 pcp1-HA-hphMX6 cut7-446* |
|  | DI 486 | *h- leu1-32 ura4-D18 lys1::P<sup>atb2</sup>-GFP-FLAG-6xHis-ura4+ sid4-tdtomato-natMX6 pcp1-HA-hphMX6 cut7-446* |
|  | DI 7 | *h- leu1-32 ura4-D18* |
| Figure 3 | DI 492 | *h- leu1-32 ura4-D18 lys1::P<sup>atb2</sup>-DmSAS6-GFP-FLAG-6xHis-ura4+ sid4-tdtomato-natMX6 pcp1-HA-hphMX6 cut7-446* |
|  | DI 631 | *h- leu1-32 ura4-D18 lys1::P<sup>atb2</sup>-DmSAS6-GFP-FLAG-6xHis-ura4+ sid4-tdtomato-natMX6 pcp1-14-HA-hphMX6 cut7-446* |
|  | DI 706 | *h+ leu1-32 ura4-D18 lys1::P<sup>atb2</sup>-DmBld10-GFP-FLAG-6xHis-ura4+ sid4-tdtomato-natMX6 pcp1-HA-hphMX6 cut7-446* |
|  | DI 710 | *h+ leu1-32 ura4-D18 lys1::P<sup>atb2</sup>-DmBld10-GFP-FLAG-6xHis-ura4+ sid4-tdtomato-natMX6 pcp1-14-HA-hphMX6 cut7-446* |
|  | DI 709 | *h+ leu1-32 ura4-D18 lys1::P<sup>atb2</sup>-6xHis-FLAG-YFP-DmSAS4-ura4+ sid4-tdtomato-natMX6 pcp1-HA-hphMX6 cut7-446* |
|  | DI 719 | *h- leu1-32 ura4-D18 lys1::P<sup>atb2</sup>-6xHis-FLAG-YFP-DmSAS4-ura4+ sid4-tdtomato-natMX6 pcp1-14-HA-hphMX6 cut7-446* |
|  | DI 7 | *h- leu1-32 ura4-D18* |
| Figure 4B | DI 105 | *h+ leu1-32 ura4-D18 lys1::P<sup>atb2</sup>-DmSAS6-GFP-FLAG-6xHis-ura4+* |
| Figure 4C, D | DI 636 | *h- leu1-32 ura4-D18 lys1::P<sup>atb2</sup>-DmSAS6-GFP-FLAG-6xHis-ura4+ sfi1-CFP-natMX6* |
|  | DI 638 | *h- leu1-32 ura4-D18 lys1::P<sup>atb2</sup>-GFP-FLAG-6xHis-ura4+ sfi1-CFP-natMX6* |
| Figure 4D | DI 636 | *h- leu1-32 ura4-D18 lys1::P<sup>atb2</sup>-DmSAS6-GFP-FLAG-6xHis-ura4+ sfi1-CFP-natMX6* |
|  | DI 638 | *h- leu1-32 ura4-D18 lys1::P<sup>atb2</sup>-GFP-FLAG-6xHis-ura4+ sfi1-CFP-natMX6* |
| Figure S1A, B | DI 456 | *h- leu1-32 ura4-D18 lys1::P<sup>atb2</sup>-DmSAS6-GFP-FLAG-6xHis-ura4+ sid4-tdTomato-natMX6* |
|  | DI 190 | *h- leu1-32 ura4-D18 lys1::P<sup>atb2</sup>-DmBld10-GFP-FLAG-6xHis-ura4+ sid4-mRFP-natMX6* |
|  | DI 202 | *h- leu1-32 ura4-D18 lys1::P<sup>atb2</sup>-6xHis-FLAG-YFP-DmSAS4-ura4+ sid4-mRFP-natMX6* |
|  | DI 655 | *h- ura4-D18 lys1::nmt1-6xHis-FLAG-YFP-DmAna2-ura4+ sid4-tdtomato-natMX6* |
|  | DI 665 | *h+ ura4-D18 leu1::nmt41-6xHis-FLAG-GFP-DmPlk4-ura4+ sid4-tdtomato-natMX6* |
|  | DI 7 | *h- leu1-32 ura4-D18* |

**Table S3.**
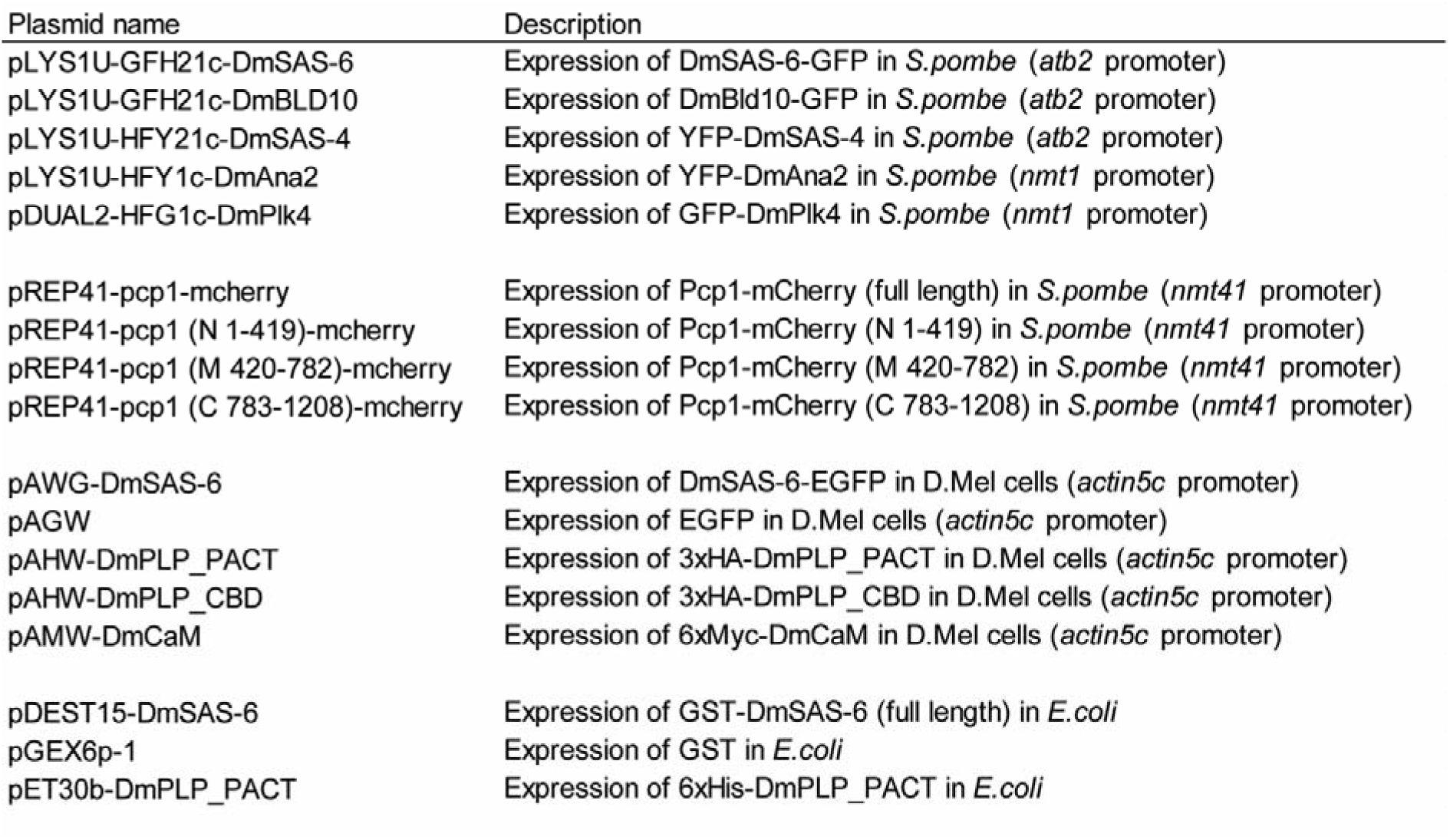
Plasmids used in this study

**Table S4.**
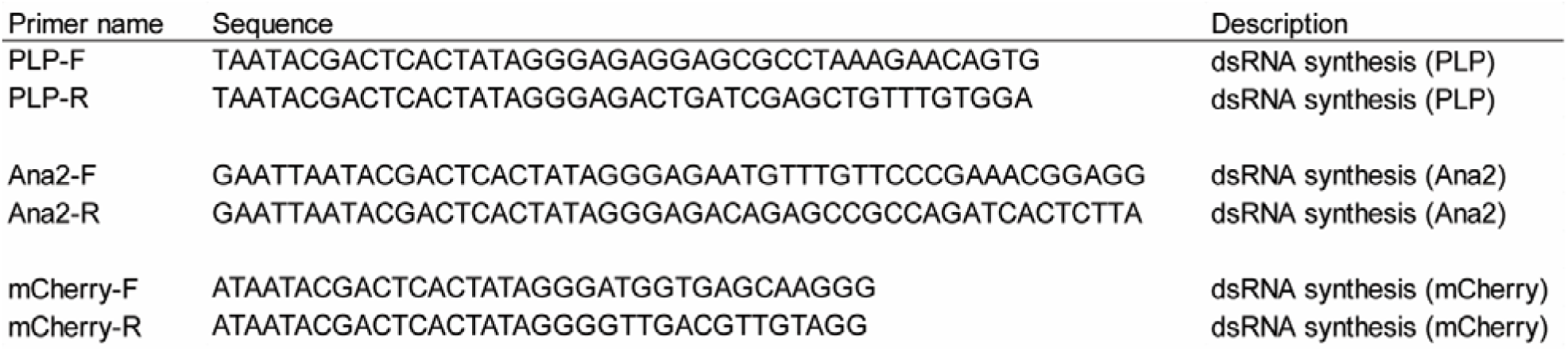
Primers used to amplify DNA templates for dsRNA synthesis

**Table S5.**
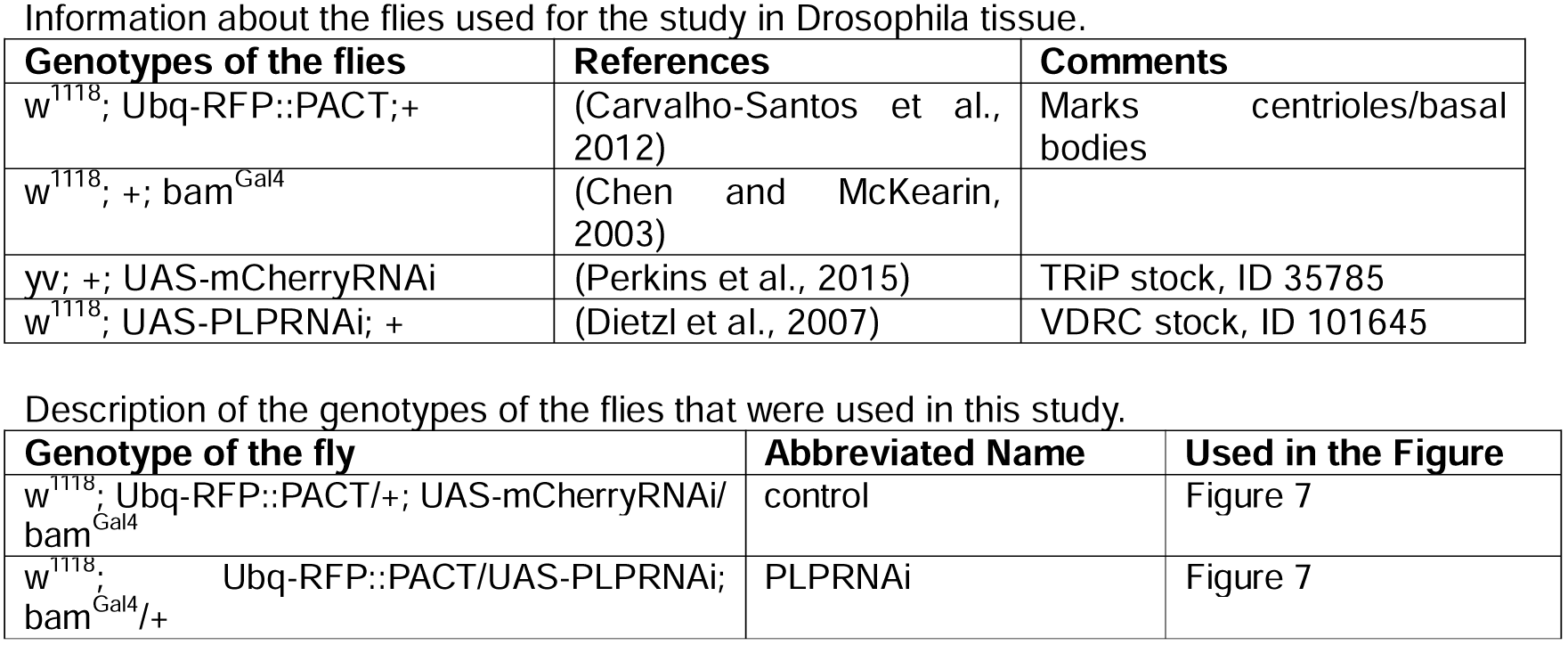
Flies used in this study

